# Prodomain-driven enzyme dimerization, a potent pH-dependent autoinhibition mechanism to control *Plasmodium* Sub1 subtilisin-like activity prior to prime merozoite egress

**DOI:** 10.1101/2023.07.31.551325

**Authors:** Mariano Martinez, Anthony Bouillon, Sébastien Brule, Bertrand Raynal, Ahmed Haouz, Pedro M. Alzari, Jean-Christophe Barale

## Abstract

Malaria symptoms are associated with the asexual multiplication of *Plasmodium falciparum* within human red blood cells (RBC) and fever peaks coincide with the egress of daughter merozoites following the rupture of the parasitophorous vacuole (PV) and the RBC membranes (RBC). Over the last two decades it has emerged that the release of competent merozoites is tightly regulated by a complex cascade of events, including the unusual multi-step activation mechanism of the pivotal subtilisin-like protease 1 (Sub1) that takes place in three different cellular compartments and remains poorly understood. Following an initial auto-maturation in the Endoplasmic Reticulum (ER) between its pro- and catalytic domains, the Sub1 prodomain (PD) undergo further cleavages by the parasite aspartic protease plasmepsin X (PmX) within acidic secretory organelles that ultimately lead to full Sub1 activation upon discharge into the parasitophorous vacuole (PV). Here we report the crystal structure of full-length *P. falciparum* Sub1 (PfS1_FL_) and demonstrate, through structural, biochemical and biophysical studies, that the atypical *Plasmodium-*specific Sub1 PD directly triggers the assembly of inactive enzyme homodimers at acidic pH, whereas Sub1 is primarily monomeric at neutral pH. Our results shed new light into the finely tuned Sub1 spatiotemporal activation during secretion, particularly the different compartmentalization of PmX processing and full Sub1 activation, and uncover a robust mechanism of pH-dependent subtilisin autoinhibition involved in the tight regulation of *P. falciparum* merozoites egress from host infected cells.

**Significance:** Malaria fever spikes are due to the rupture of infected erythrocytes, allowing the egress of *Plasmodium* sp. merozoites and further parasite propagation. This fleeting tightly regulated event involves a cascade of enzymes, culminating with the complex activation of the subtilisin-like protease 1, Sub1. Differently than other subtilisins, Sub1 activation strictly depends upon the processing by a parasite aspartic protease within acidic merozoite secretory organelles. However, Sub1 biological activity is requested in the pH neutral parasitophorous vacuole, to prime effectors involved in the rupture of the vacuole and erythrocytic membranes. Here we show that the unusual, parasite specific Sub1 prodomain is directly responsible for its acidic- dependent dimerization and autoinhibition, required for protein secretion, before its full activation at neutral pH in a monomeric form. pH-dependent Sub1 dimerization defines a novel, essential regulatory element involved in the finely tuned spatiotemporal activation of the egress of competent *Plasmodium* merozoites.

## Introduction

Subtilisins, or subtilases, compose a very diverse family of ubiquitous serine proteases involved in a broad spectrum of biological functions (1–3). According to the MEROPS database (http://merops.sanger.ac.uk, 4), subtilases constitute the S8 family in clan SB of serine proteases and are divided in the S8A and S8B sub-families corresponding respectively to prokaryotic enzymes (1, 2), known to display a broad specificity, and to eukaryotic kexin-like enzymes (3). Among the latter, the Ca^2+^-dependent prohormone convertases (PCs) precisely mature polypeptidic precusors after mono or di-basic residues, producing functionnal hormones, also activating bacterial toxins or surface proteins of various viruses or bacterial toxins (5–7). As most other proteases, subtilisins are synthetized as an inactive zymogen with a core structure composed of at least two domains, the catalytic domain and a dual function prodomain. The catalytic domain, called the peptidase S8 domain, is organized around a catalytic His-Asp-Ser triad, with a fourth conserved Asn that is part of the oxy-anion hole that stabilizes the negative charge of the tetrahedral intermediate formed between the enzyme and its substrate (8). Subtilisin prodomains act both as intramolecular chaperones of and potent inhibitors of their cognate catalytic domains (9–12). In both bacteria and eukaryotic cells, primary automaturation occurs between the subtilase catalytic domain and its prodomain during their secretion, when exposed to variations of pH and/or Ca^2+^-concentration that induce complex disassembly and subsequent degradation of the prodomain by the active enzyme (12, 13).

To avoid collateral damage due to inappropriate protease activation, different regulatory mechanisms exist to ensure that the enzyme activation is triggered at the right place and time. Thus, in bacteria the removal of subtilisin prodomains usually depends upon two distinct autoproteolytic cleavages, each with a different pH optimum (14, 15). In the case of the PCs, specific histidines located in prodomains have been shown to act as pH sensors to trigger subtle conformational changes in the desired cellular compartment to release and degrade their prodomain. Furin His_69_ and PC1/3 His_72_/His_75_ have been shown to play this role during their secretion in the mildly acidic pH of the trans-Golgi network (pH ::6.5) and in more acidic mature secretory granules (pH ::5.5), respectively (16, 17). While pH also plays an important role to remove the prodomain of plant subtilisins (18), these enzymes also frequently contain, between the histidine and the serine of their catalytic triad, a protease associated (PA) domain described to trigger protein-protein interactions (19). While involved in extended substrate recognition (20), the PA domain also regulates the activity of the SlSBT3 plant subtilisin, promoting its vital homodimerization to maintain the active site accessible to the substrate (21, 22). Finally, subtilisins may also be regulated in *trans*, by prodomain-like polypeptides that inhibit cellular enzymes found in yeast (23), plant (24) or in the apicomplexa human-infecting parasites *Toxoplasma gondii* (25, 26) or *Plasmodium falciparum* (27).

The subtilisin-like protease Sub1 plays an essential role at different stages of *Plasmodium* sp., the etiological agents of malaria, the most devastating parasitic disease which control is compromised by the constant selection of multi-resistances to existing treatments (28). Transmitted through the bite of the *Anopheles* mosquito vector, the parasites quickly reach the liver where they invade host hepatocytes in which they asexually multiply in merozoites within a parasitophorous vacuole (PV). Following this asymptomatic phase, Sub1 plays a critical role allowing the egress of liver merozoites (29, 30) that reach the blood where they invade host red blood cells (RBC) to initiate the erythrocytic cycle responsible for the symptoms of malaria. As in the hepatocytes, the parasites grow and asexually multiply by schizogony within a parasitophorous vacuole in the erythrocyte cytoplasm in 24 to 72 hours, depending on *Plasmodium* species. The newly formed merozoites egress after the rupture of the PV and erythrocytic membranes before to invade new RBC. The egress of merozoites is a finely regulated and multi-step event initiated by a kinases-mediated (31, 32) intracellular shift of calcium that promotes the discharge of Sub1 from a merozoite secretory organelle, the exoneme, to the PV (33). Once there, active Sub1 processes and activates soluble proteases of the SERA family involved in the rupture of the PV membrane, and matures different merozoite surface proteins involved in the invasion of RBC (34–36). Despite its central role in the cascade of enzymes that controls the egress of mature merozoites (37, 38), important mechanistic questions on Sub1 transport and activation remain unknown.

*Plasmodium* sp. Sub1 are synthesised as inactive precursors of ::80 kDa with a bacterial-like C-terminal catalytic domain (39, 40) and an atypical N-terminal prodomain (PD) composed of two regions: a classical bacterial subtilisin-like prodomain preceded by a parasite-specific N-terminal insertion of unknown function. Following translocation into the endoplasmic reticulum (ER), Sub1 undergoes autocatalytic cleavage to yield a C-terminal 54- kDa intermediate (p54) that remains noncovalently bound to the cleaved PD of 31 kDa (p31). During exoneme transport to the PV, Sub1 undergoes further processing by the Asp protease plasmepsin X (PmX) (41–43), which has recently been shown to directly cleave both the Sub1 PD and the catalytic domain (44). However, it remains unclear why PD processing within the exoneme is spatially dissociated from full Sub1 activation, which only occurs at a later stage, upon discharge into the PV. Here we report the crystal structure of full-length *P. falciparum* Sub1 (PfS1_FL_) and demonstrate, through comparative structural analysis substantiated by biochemical and biophysical studies, that the atypical Sub1 PD is directly responsible for the assembly of inactive enzyme homodimers at acidic pH. Our results uncover a novel robust mechanism of pH-dependent subtilisin autoinhibition, which plays a crucial role in the tight regulation of *Plasmodium* merozoite egress from host cells and can account for the different compartmentalization of PmX processing and full Sub1 activation.

## Results

### The overall structure of full-length PfS1

The X-ray structure of recombinant full-length protease Sub1 from *P. falciparum* (PfS1_FL_) expressed in insect cells was determined by molecular replacement methods at 3.09 Å resolution (Table 1). As expected, the structure revealed a subtilisin-like catalytic core (residues 330-668) tightly bound to its cognate PD composed of two distinct subunits (residues 37-97 and 137-217, Figure 1a). Complex formation between the catalytic core and the PD buries ∼2,350 Å^2^ of surface area from each moiety, which roughtly duplicates the values observed for the similar interface in bacterial subtilisin (45). The overall architecture is very similar to that of the closely related Sub1 from *Plasmodium vivax* (PvS1_FL_ (46), PDB code 4tr2) (Figure S1), as the two structures can be superimposed with an overall rmsd of 0.78 Å for 462 equivalent Cα atomic positions. All structural features characteristic of *Plasmodium* subtilases are conserved, including the three high-affinity calcium-binding sites in the catalytic domain, two of which are specific to *Plasmodium* subtilases, and a fourth calcium-binding site in the PD, all of which were previously described for PvS1_FL_ (46). Another common feature is that the PD had undergone autocleavage at its primary maturation site (Asp 217 in PfS1_FL_ and Asp202 in PvS1_FL_) but remained tightly associated to the catalytic core in the crystal.

**Figure 1.**
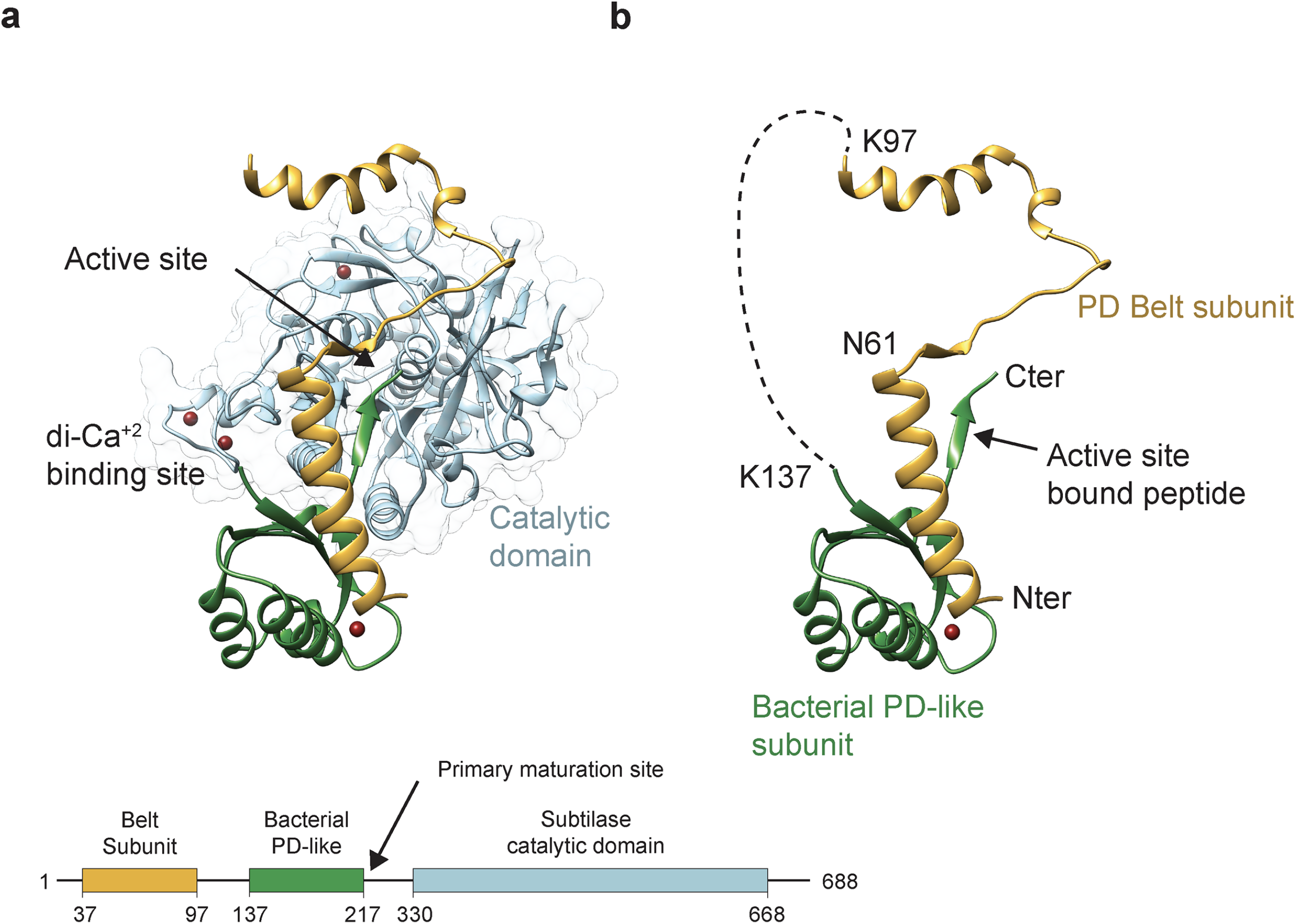
Overall structure of full-length *P. falciparum* Sub1 (PfS1FL). (a) The catalytic domain (in cyan) is tightly bound to its cognate PD made up of two distinct subunits, the N- terminal belt subunit (yellow) and the bacterial PD-like subunit (green). The PD connecting segment between the two subunits (residues 116-153) and the N-terminal region of the mature catalytic domain (residues 236-347) are not visible in the electron density and are presumably disordered in the crystals. The schematic domain organization of PfS1_FL_, with the three structured domains, is shown below the structure. **(b)** Detailed structure of the Sub1 PD colour-coded as in panel (a). The active site-bound propeptide and specific residues discussed in the text are labelled.

**Table 1.**
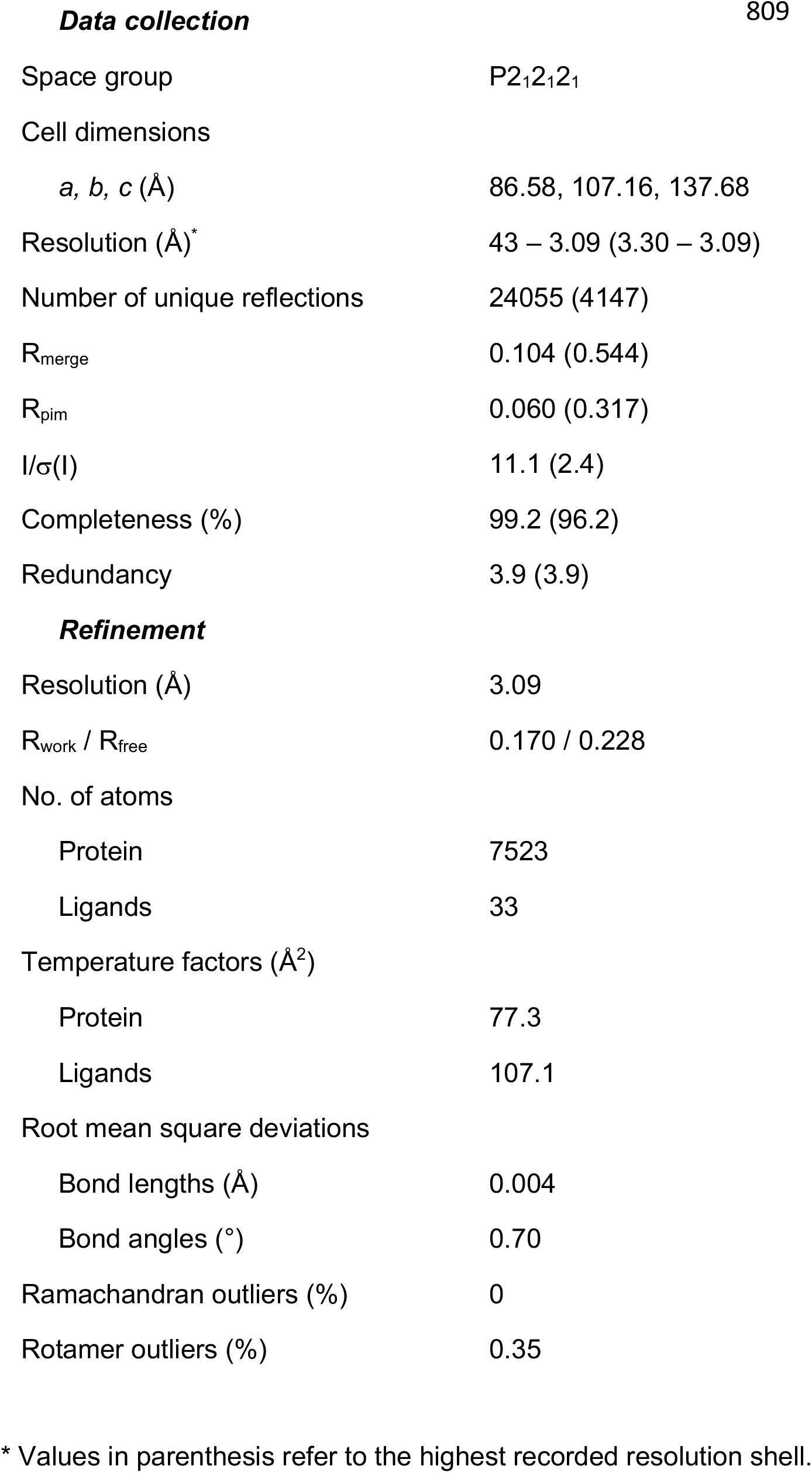
Crystallographic data.

|  |  |
| --- | --- |
| Space group | P2<sub>1</sub>2<sub>1</sub>2<sub>1</sub> |
| Cell dimensions |  |
| *a*, *b*, *c* (Å) | 86.58, 107.16, 137.68 |
| Resolution (Å)* | 43 – 3.09 (3.30 – 3.09) |
| Number of unique reflections | 24055 (4147) |
| R<sub>merge</sub> | 0.104 (0.544) |
| R<sub>pim</sub> | 0.060 (0.317) |
| I/σ(I) | 11.1 (2.4) |
| Completeness (%) | 99.2 (96.2) |
| Redundancy | 3.9 (3.9) |
| ***Refinement*** |  |
| Resolution (Å) | 3.09 |
| R<sub>work</sub> / R<sub>free</sub> | 0.170 / 0.228 |
| No. of atoms |  |
| Protein | 7523 |
| Ligands | 33 |
| Temperature factors (Å<sup>2</sup>) |  |
| Protein | 77.3 |
| Ligands | 107.1 |
| Root mean square deviations |  |
| Bond lengths (Å) | 0.004 |
| Bond angles (°) | 0.70 |
| Ramachandran outliers (%) | 0 |
| Rotamer outliers (%) | 0.35 |

\* Values in parenthesis refer to the highest recorded resolution shell.

The Sub1 PD is made up of two distinct subunits and displays an atypical folding (Figure 1b). The structural core consists of an N-terminal α-helix that interacts through an interface of 700 Å^2^ with an α/β subunit similar to bacterial subtilisin PDs (45), which includes the C-terminal peptide extension that remains bound to the protease active site after primary maturation. The N-terminal helix is connected to the first β-strand by a 75 residues-long loop that is largely devoid of secondary structure and mostly disordered, except for its first part that embraces the catalytic domain as a belt (Figure 1a). The first structural subunit of the PD (residues 37- 97, referred to here as the belt subunit) includes the *Plasmodium*-specific N-terminal helix and the ordered part of the adjacent loop. The belt subunit is also present in PvS1_FL_ and is highly conserved in all *Plasmodium* species (46), suggesting that it fulfills an important, as yet unknown, functional role. This subunit was missing in the previously reported structure of *P. falciparum* Sub1 in complex with a specific antibody (PDB code 4lvn) (47), possibly due to N-terminal truncation of the protein following chymotrypsin treatment during purification.

### The Sub1 prodomain mediates pH-dependent protein dimerization

Sub1 is active as a monomer at neutral pH. However, PfS1_FL_ crystallized as a homodimer at acidic pH, with an interfacial area of ∼1600 Å^2^ that involves extensive intermolecular hydrophobic and H-bond interactions. The PD is largely responsible for fastening together the Sub1 homodimer, as the belt subunits from both protomers occupy the center of the dimer, sandwiched between the two catalytic domains with their active sites facing each other (Figure 2a). Intermolecular interactions between the belt subunits from the two protomers (through an interface of 1050 Å^2^) accounts for roughly two thirds of the total dimer interface (Figure 2b). Other intermolecular contacts involve the interactions between the catalytic domain from one protomer with the belt subunit from the opposite protomer (250 Å^2^ interface). In contrast, the two catalytic domains are not directly in contact with each other, except for the tip of the prominent loop C521-C534, which protrudes out from one monomer to clamp the opposite catalytic domain (Figure 2a). This loop, which is conserved in *Plasmodium* Sub1 orthologs but absent from other subtilisins, is stabilized by a disulphide bridge (Cys521-Cys534) at its basis and is adjacent to the oxyanion hole residue Asn520. Previous work had suggested that this redox-sensitive disulphide could act as a regulator of protease activity in the parasite (47).

**Figure 2.**
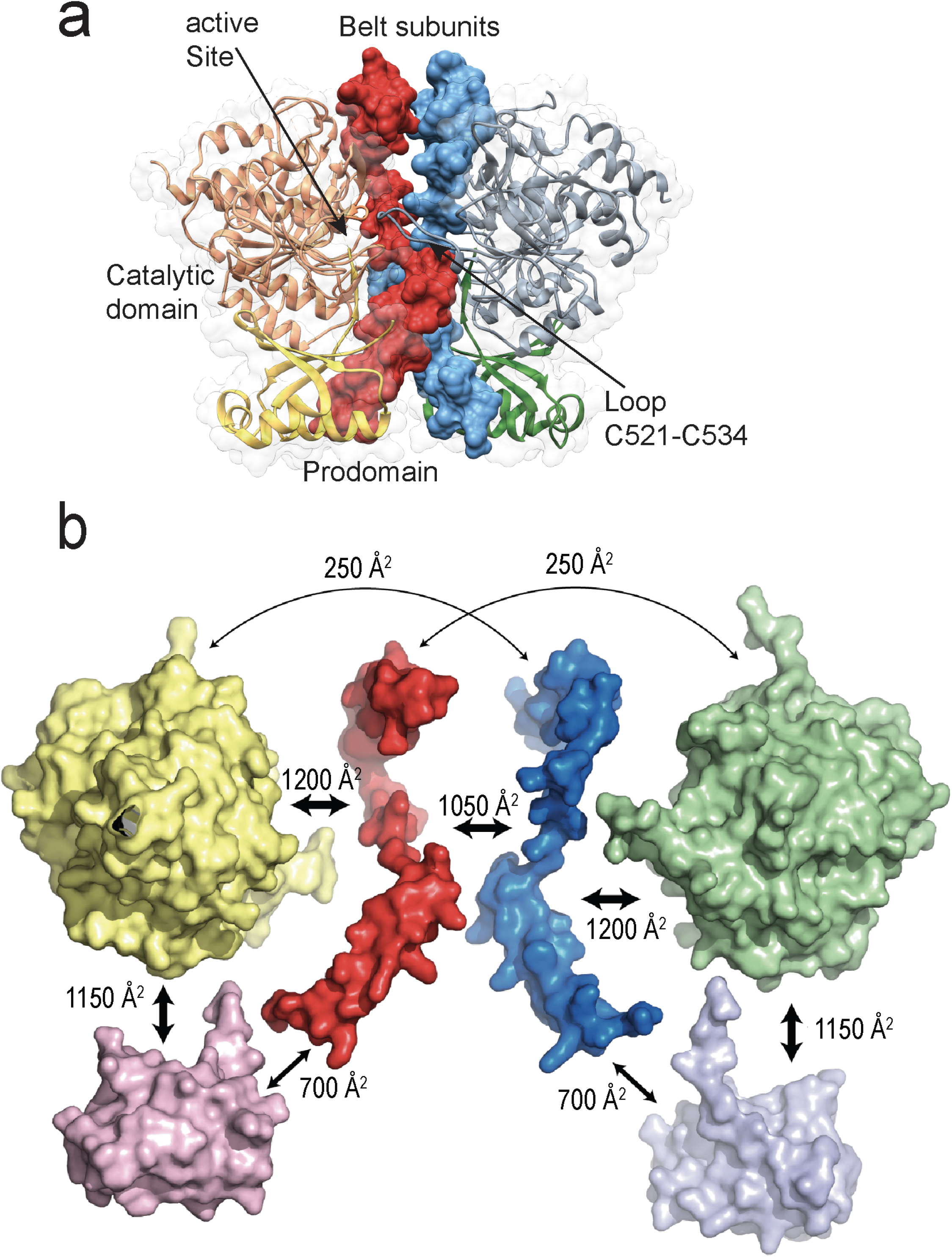
The *Plasmodium*-specific belt subunit mediates Sub1 dimerization. (a) Crystallographic dimer of PfS1_FL_ showing the interactions beyween the two protomers. The belt subunits (shown in red and blue with molecular surface representation) occupy the center of the dimer interface. **(b)** Exploded view of the PfS1_FL_ homodimer showing the interface contact surfaces between the different subdomains. The catalytic domains are shown in yellow and green, the bacterial PD-like subunits in pink and light blue, and the belt subunits in red and blue. Due to the presence of the belt subunit, the total contact surface between the PD and the catalytic domain within a single Sub1 monomer (3350 Å^2^) roughly duplicates the interface observed in other subtilisins.

Interestingly, a similar face-to-face dimer (rmsd of 1.58 Å^2^ for 920 equivalent residues) had been observed in crystals of PvS1_FL_ (Figure S2a), which were also grown in acidic conditions (pH 4.2 - 5) but belong to a different crystal form (46). In PvS1_FL_, however, a different crystal lattice interface (Figure S2b) presented a larger contact surface area (∼1800 Å^2^ compared to ∼1400 Å^2^ for the face-to-face dimer). To experimentally investigate whether any of these interactions could stabilize an oligomeric form of Sub1 in solution, we carried out analytical ultracentrifugation studies (AUC) of recombinant PvS1_FL_. The protein was found to form a stable homodimer in solution at pH 5 but behaves as a monomer at pH 8 under otherwise identical conditions (Figure 3a and Table 2). These results imply that (i) Sub1 dimerization is pH-dependent, as an increase of pH promoted dimer dissociation, and (ii) one of the two crystallographic dimers observed for PvS1_FL_ (Figure S2) should be stable in solution. To elucidate which one, we repeated the AUC experiment on a N-terminal truncated form of PvS1 lacking the belt subunit (PvS1β_Belt_), as it is largely involved in the face-to-face dimeric interface but not in the back-to-back dimer (Figure S2). Deletion of the N-terminus rendered the protein monomeric in solution at acidic pH (Figure 3b and Table 2), indicating that the face-to-face dimer observed for both the PvS1_FL_ and PfS1_FL_ structures in completely different crystal environments does correspond to a stable Sub1 dimer in solution.

**Figure 3.**
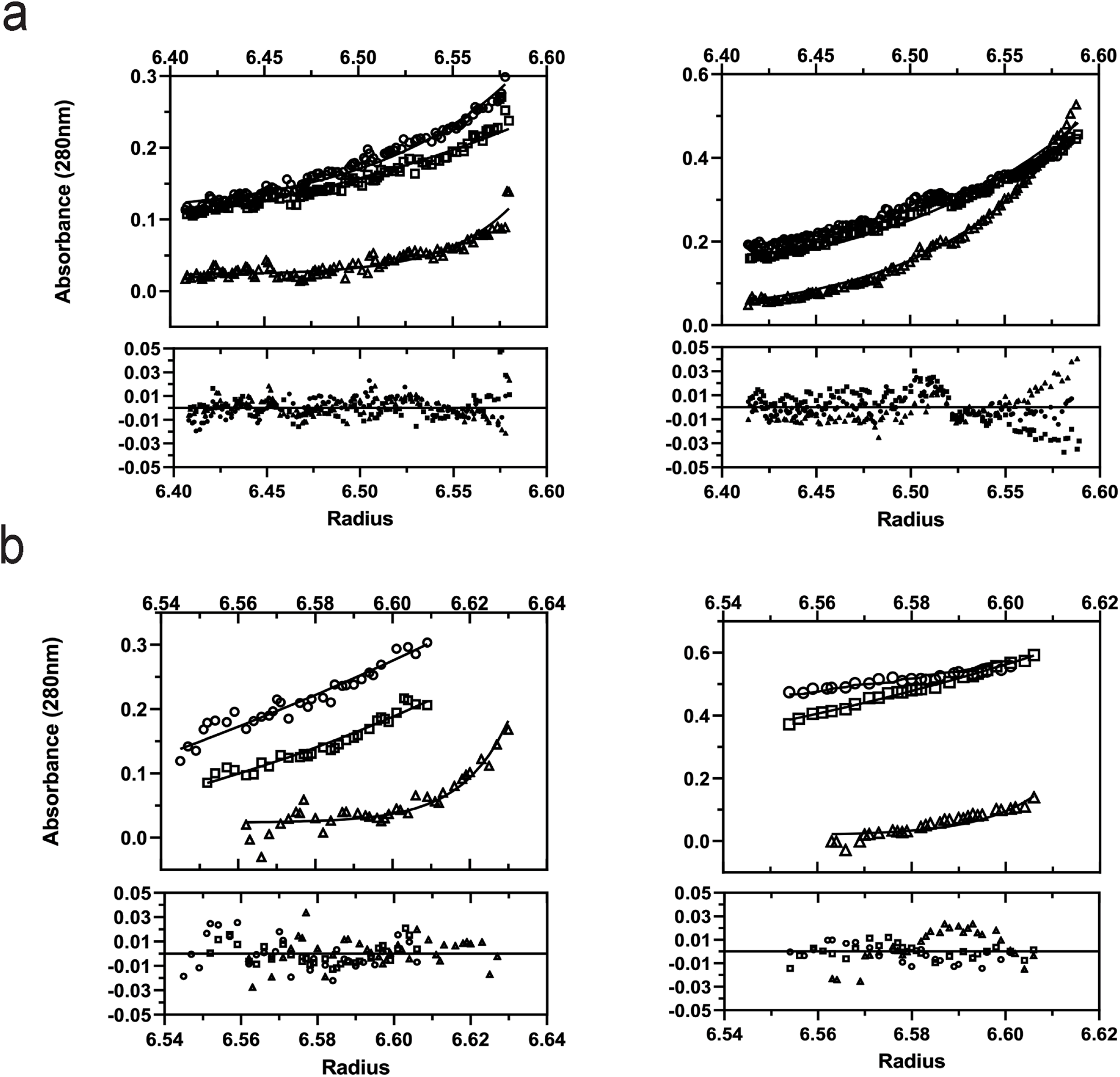
pH-dependent dimerization of Sub1. (a) Analytical ultracentrifugation (AUC) sedimentation equilibium analysis on purified PvS1_FL_ at pH 5 (left panel) and pH 8 (right panel). AUC profiles were recorded at 20 °C using different concentrations at three rotor speeds 9000 rpm (squares), 13000 rpm (circles) and 16000 rpm (triangles). For clarity, only equilibirum traces of proteins at 5 µM are shown. For every experiment, the upper panel shows the sedimentation equilibrium profiles with the lines of best fit obtained for a single species model (molecular weights are shown in Table 2), and the lower panel shows the data fitting residuals. **(b)** Similar AUC sedimentation analysis on purified PvS1τι_Belt_ at pH 5 (left panel) and pH 8 (right panel). As above, the best fit was obtained for a single species model (Table 2).

**Table 2.** pH-dependent oligomerization of PvS1_FL_ and PvS1ι1_Belt_.

| Protein | Theor. MW<br>(monomer) | pH 5 | pH 8 |
| --- | --- | --- | --- |
| PvS1<sub>FL</sub> | 67.71 kDa | 135.8 ± 2.8 kDa | 67.9 ± 1.1 kDa |
| PvS1<sub>ΔBelt</sub> | 56.57 kDa | 57.5 ± 0.2 kDa | 57.5 ± 4.5 kDa |

At neutral pH, the entire dimerization interface is predicted to be negatively charged (Figure 4a), which likely results in Coulombic repulsion and prevents dimerization. Largely buried at the center of the interface, the belt subunit contains an important concentration of negatively charged residues (Figure 4b). In PfS1_FL_, 18 out of 61 ordered residues in this region (29.5%) are Asp or Glu, whereas in the rest of the structure this percentage is significantly lower, 12.4%, matching the average amino acid protein composition in whole proteomes (48). Importantly, most of these negatively charged belt positions are largely or strictly conserved in *Plasmodium* Sub1 orthologs (Figure 4c), and some of them are partially occluded from the solvent upon dimerization. At pH 5 the dimerization interface of PfS1 is predicted to become neutral (Figure 4a), thus favoring dimerization. Several negatively charged residues that belong to or interact with the belt subunit are predicted to become protonated at pH 5 using the PROPKA program (49). For example, a cluster of seven glutamic acids (E38, E46, E51, E187, E202, E209, and E495), five of which would be protonated at pH 5, participate in the interaction of the N-terminal belt helix with the bacterial-like pro-region and the catalytic domain from the same molecule (Figure S3). The close proximity between their carboxylate groups suggest that Coulombic repulsion at neutral pH would destabilize this association. Moreover, eight histidines are predicted to change from neutral to positively charged at pH 5, three of which (H464, H523 and H596) are part of the homodimer interface. These pH-induced charge transitions might allow the formation of intermolecular salt bridges at acidic pH, as for example H464 and H596 from each protomer respectively face D68 and E71 from the other protomer in the dimer. In summary, these changes in surface charge can explain the observed Sub1 transition in solution from a single dimeric species at acidic pH to a single monomeric form at neutral pH.

**Figure 4.**
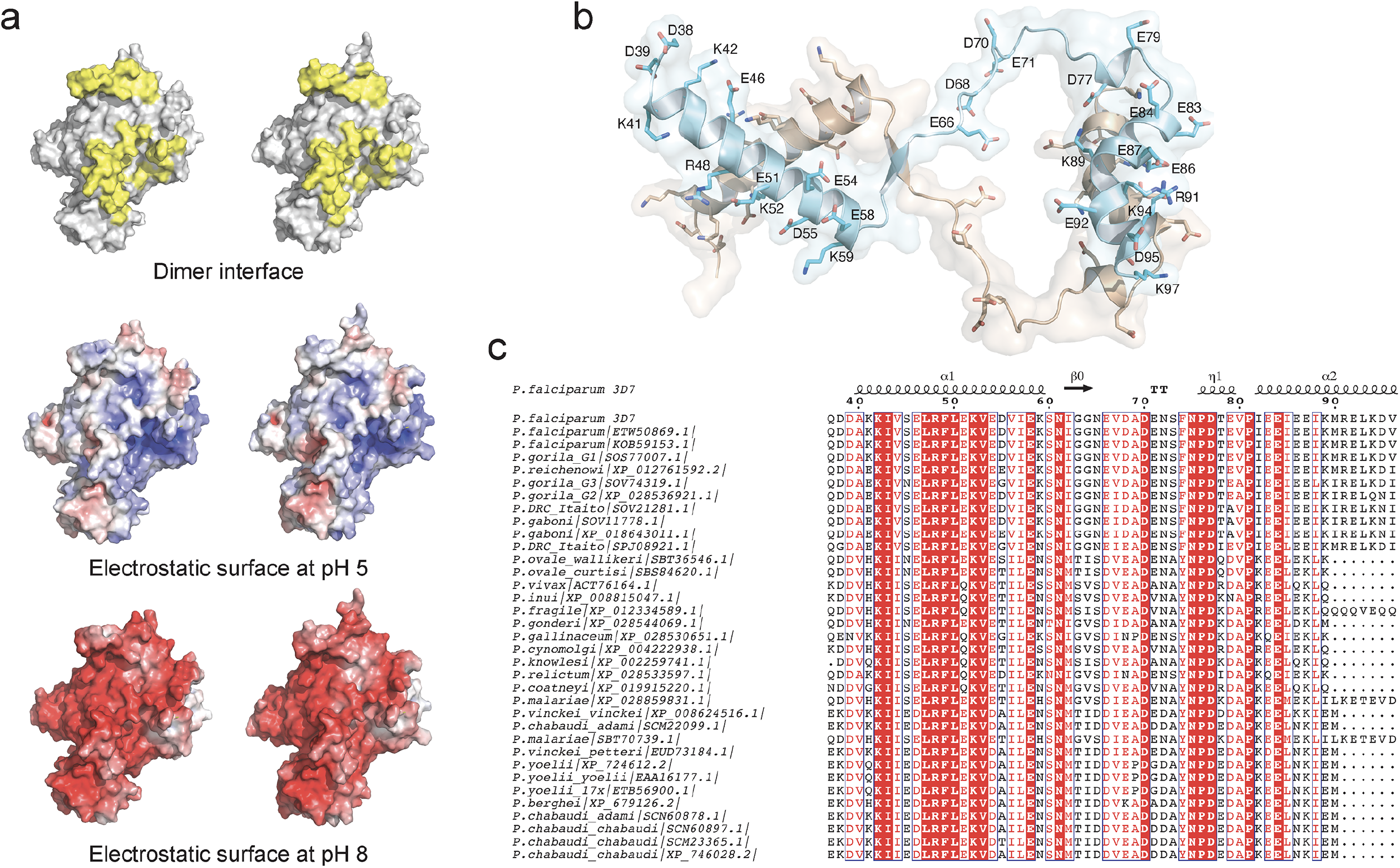
pH-dependent interactions. (a) Open book view of the dimer in surface representation (top) with the interface residues colored yellow. Below, gradient visualization from red (-10 kT/e) to blue (+10 kT/e) of the electrostatic surface at pH 5 and 8, calculated with the APBS server (75, 76) based on residue pKa determined by PROPKA (49). **(b)** Cartoon representation of the belt subunits from both protomers, which occupy the center of the homodimer. In this orientation, the catalytic domains (omitted from the figure) are respectively below and above the plane of the paper. All charged residues (Asp, Glu, Arg, Lys) are shown in stick representation and labeled for one of the two subunits. **(c)** Conserved amino acid sequence of the belt subunit from different *Plasmodium* Sub1 orthologs.

### Protein dimerization as a robust mechanism for enzyme autoinhibition

PD-mediated protein dimerization traps the cleaved propeptide within the active site and completely blocks the access to (or release from) the substrate-binding clefts from both protomers (Figure 5a), indicating that homodimerization could serve as an efficient mechanism for Sub1 autoinhibition. Enzymatic assays at different pH values tend to confirm this hypothesis. Recombinant PfS1 was reported to be active in a wide range of pH values, with optimal activity around pH 8 (50) and we obtained similar results for PvS1. To investigate the inhibitory effects of the PD structural subunits on PvS1 activity at different pH values, we carried out enzyme assays on the full-length enzyme (PvS1_FL_), a construct lacking the belt subunit but containing the bacterial PD-like subunit (PvS1β_Belt_) and the catalytic domain alone (PvS1_CD_). For all the constructs, the enzymatic activity could be detected in a wide range of tested pH values, with a maximum around pH 7 (Figure 5b). As expected, PvS1_CD_, which corresponds to the full active protease, has a significantly higher activity than the two other constructs, which are partially or totally inhibited by the bound PD and whose activity can still be measured possibly because a fraction of the processed catalytic domains have lost their bound cognate PD during purification (50). Interestingly, no detectable activity was observed for dimeric PvS1_FL_ at the most acidic conditions (pH 5 – 5.5), whereas a small but significant activity could still be measured at the same pH for both PvS1β_Belt_ and PvS1_CD_ (both unable to dimerize), highlighting the robustness of pH-induced dimerization as an autoinhibition mechanism.

**Figure 5.**
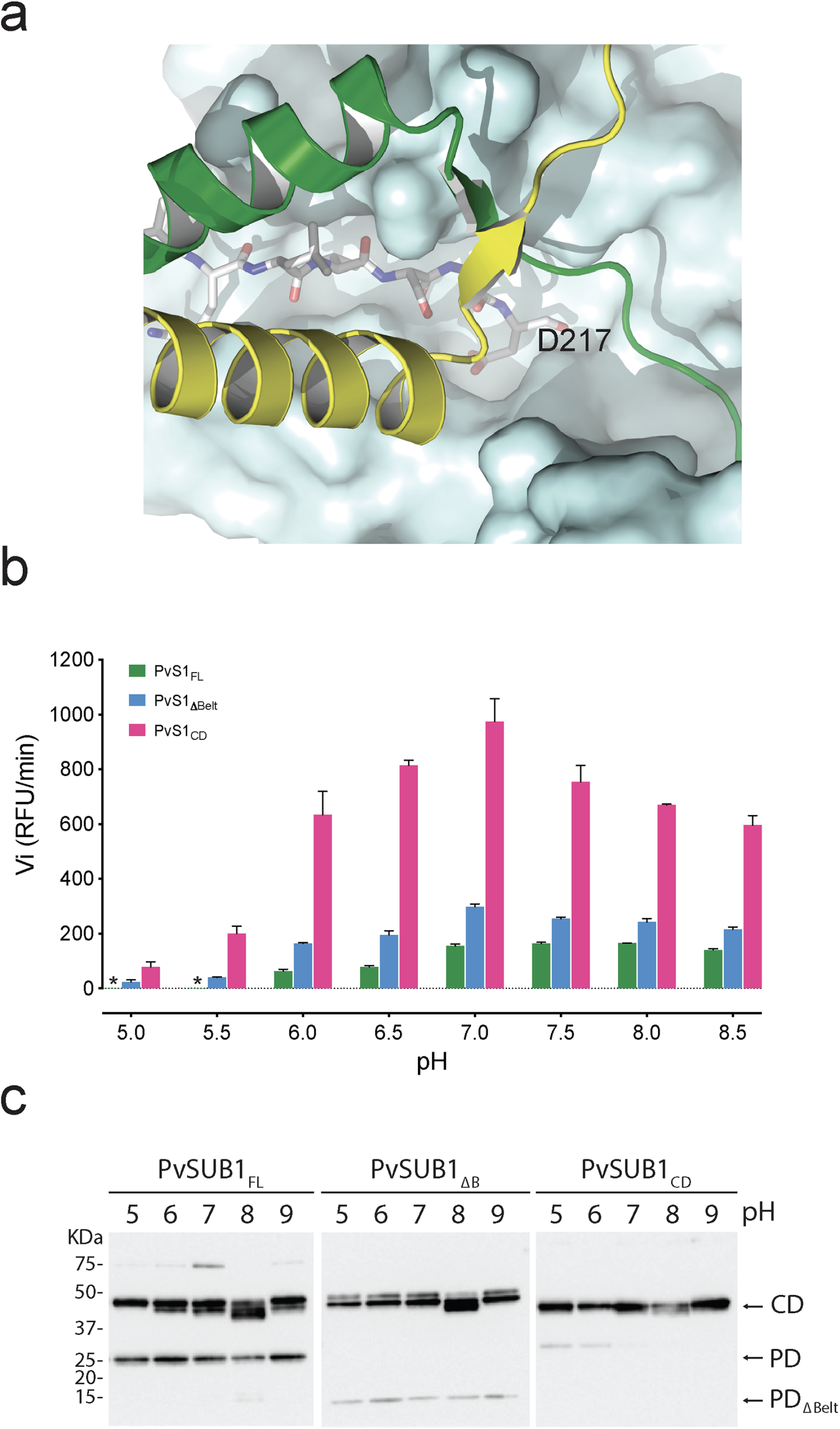
Protein dimerization as a robust inhibition mechanism . (a) The active site cleft of the catalytic domain (represented as a blue molecular surface) containing the cleaved prodomain peptide (in stick representation) is completely blocked by the two belt subunits (yellow and green) in the homodimer. The second catalytic domain (above the plane of the paper in this view) has been omitted for clarity. **(b)** pH dependence of enzyme activity for different PvS1 constructs as indicated. The initial hydrolysis rates (arbitrary units) are shown for the actual final pH of the assay buffer at 37 °C, determined as described under “Experimental Procedures.” In the plots, each data point represents the mean value of three independent assays and an asterisk (*) indicates undetectable activity. **(c)** SDS-PAGE for the three tested constructs at different pH values. The bands corresponding to the catalytic domain (CD), the full prodomain (PD) and the prodomain lacking the belt subunit (PDτι_Belt_) are indicated on the right.

### PfS1 is detected as a dimer in *P. falciparum* merozoites prior to egress from infected erythrocytes

To investigate how native PfS1 behaves within the parasites, proteins have been prepared from a culture of *P. falciparum* enriched in mature segmented schizonts containing merozoites prior their egress from infected RBC (Figure 6a, left panel) or from a culture at the time of merozoite egress, composed of both mature schizonts and free merozoites (Figure 6a, rigth panel). Following proteins migration on polyacrylamide gels using native conditions and transfer on nitrocellulose, antibodies raised against PfS1 revealed a main form with apparent molecular weigths of approximately 180 kDa (Figure 6b, lanes 2 and 3) that could correspond to the dimeric form of full-length PfS1 (Mw 77 kDa). A fainter signal at 66 kDa that could correspond to PD-free catalytic domain of PfS1 (Mw 53 kDa) is detected only in schizonts prior to merozoite egress under native conditions (Figure 6b lane 2). Western blot analysis with the same protein extracts separated by denaturating SDS-PAGE gel-electrophoresis and revealed with anti-PfS1 antibodies detect the expected full-length precursor and catalytic forms in both parasite cultures (Figure 6c, lanes 2 and 3), although the signal corresponding to PfS1 catalytic forms appeared fainter in the culture containing merozoites that have already egressed from infected RBC (Figure 6c, lane 3). As shown by immunofluorescence analysis, the signal revealed by anti-PfS1 antibodies shown a punctated labelling (Figure 6d), consistent with a location within the merozoite secretory vesicles (exoneme) from which Sub1 is secreted as an active enzyme in the parasitophorous vacuole, thus initiating merozoite egress.

**Figure 6.**
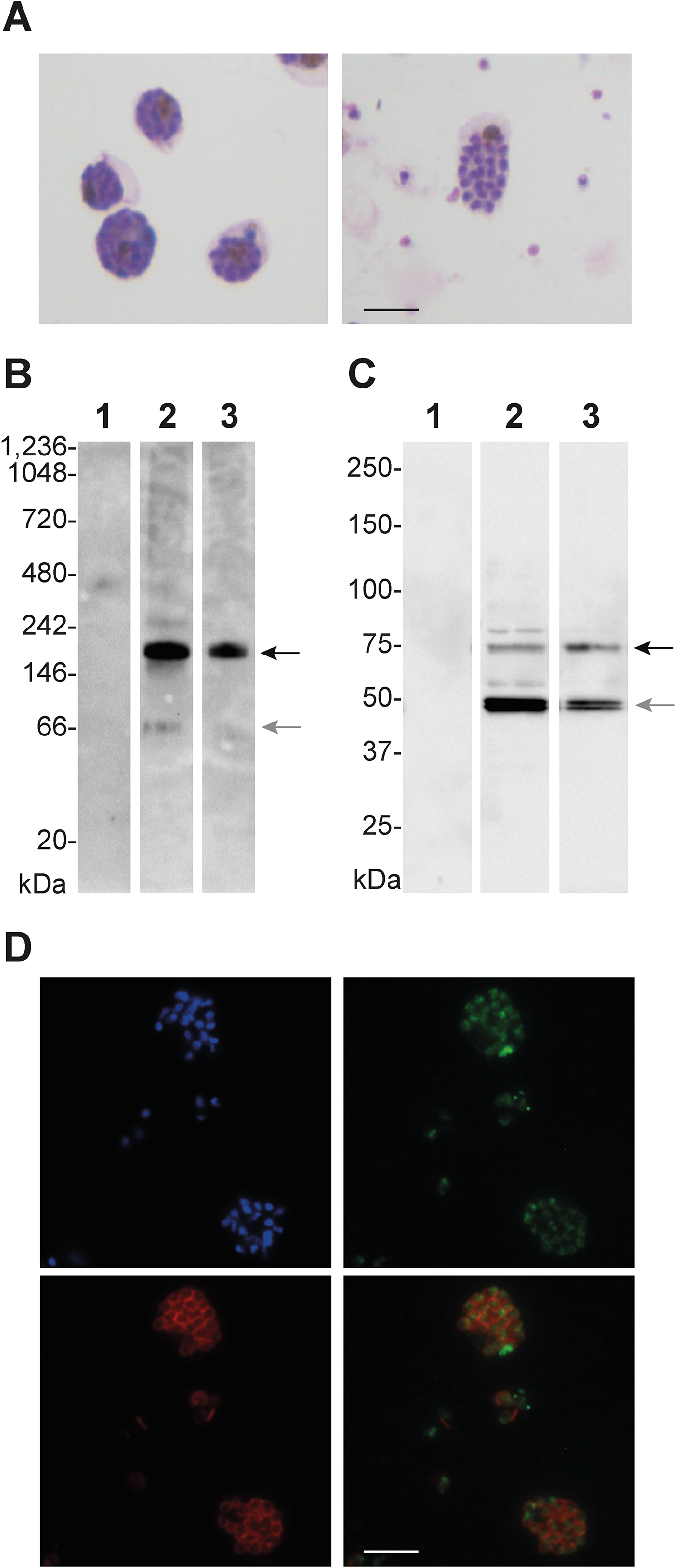
A multimeric form of PfS1 in *P. falciparum* schizonts prior to the merozoites egress. (a) Giemsa-stained parasite cultures corresponding to mature *P. falciparum* schizonts (left panel) and 5 hours later containing mature schizonts and free merozoite (rigth panel) that have egressed from infected erythrocytes. **(b)** Western blot of native protein extracts prepared from uninfected human red blood cells (Lane 1), synchronous culture of mature *P. falciparum* schizonts prior to the egress of merozoites (Lane 2, corresponding to the left panel of part A) and a mixture of mature schizonts and free merozoite (Lane 3, corresponding to the rigth panel of part A) revealed using anti-PfS1 antibodies. The multimeric form of PfS1 compatible with a dimer form of full-length PfS1 and the form compatible with PfS1 catalytic domain are indicated by black and grey arrows, respectively. Native protein molecular weights (MW) are indicated in kDa. **(c)** Western blot of boiled, reduced and migrated on denaturating SDS-PAGE electrophoresis protein extracts prepared from uninfected human red blood cells (lane 1?), synchronous culture of mature *P. falciparum* schizonts prior to the egress of merozoites (Lane 2, corresponding to the left panel of part A) and a mixture of schizonts and free merozoite (Lane 3, corresponding to the rigth panel of part A) revealed using anti-PfS1 antibodies. Traces of the precursor of full-length PfS1 and the most abundant maturated forms corresponding to PfS1 catalytic domain are indicated by black and grey arrows, respectively. Protein molecular weights (MW) are indicated in kDa. **(d)** Immunofluorescence assays using DNA-labelling of the nuclei of differenciated intra-schizonts merozoites (Hoescht, blue), anti- PfS1 antibodies (green) and antibodies raised against the Merozoite Surface Protein 1 (MSP 1, red). The bottom right panel shows an overlay of PfS1 and PfMSP1 labelling. The scale bar represents 5 µm.

## Discussion

Like other subtilisins, Sub1 is synthetized as an inactive precursor made up of a PD bound to the catalytic domain. Unlike other subtilisins, however, Sub1 is characterized by an atypical trifunctional PD capable not only of facilitating protein folding and inhibiting the protease (51), but also of inducing pH-dependent protein dimerization, as we have shown here. Primarily mediated by the belt subunit, this later function ensures a robust inhibition mechanism during Sub1 transport in the acidic environment of the exoneme, and possibly explains the essentiality of the *Plasmodium*-specific N-terminal belt extension for the growth of *P. berghei* erythrocytic stages *in vivo* (46). Taken together with previous work from different teams, our results suggest a consistent model for the activation of Sub1 in several steps (Figure 7). Initially, the inactive Sub1 precursor undergoes a first auto-catalytic cleavage at Asp217, between its PD and catalytic domain. This primary maturation takes place within the ER (52) and is required for protein dimerization, because an uncleaved propeptide within the active site would sterically clash with the position of the belt subunit in the dimer (see Figure 5a). Consistently, a mutant form of Sub1 unable to cleave itself at this primary maturation site was found to accumulate in the ER (44) and the protease was mainly found as a dimer in *P. falciparum* mature schizonts prior to merozoite egress (Figure 6), strongly suggesting that Sub1 dimerization is essential for exoneme transport.

**Figure 7.**
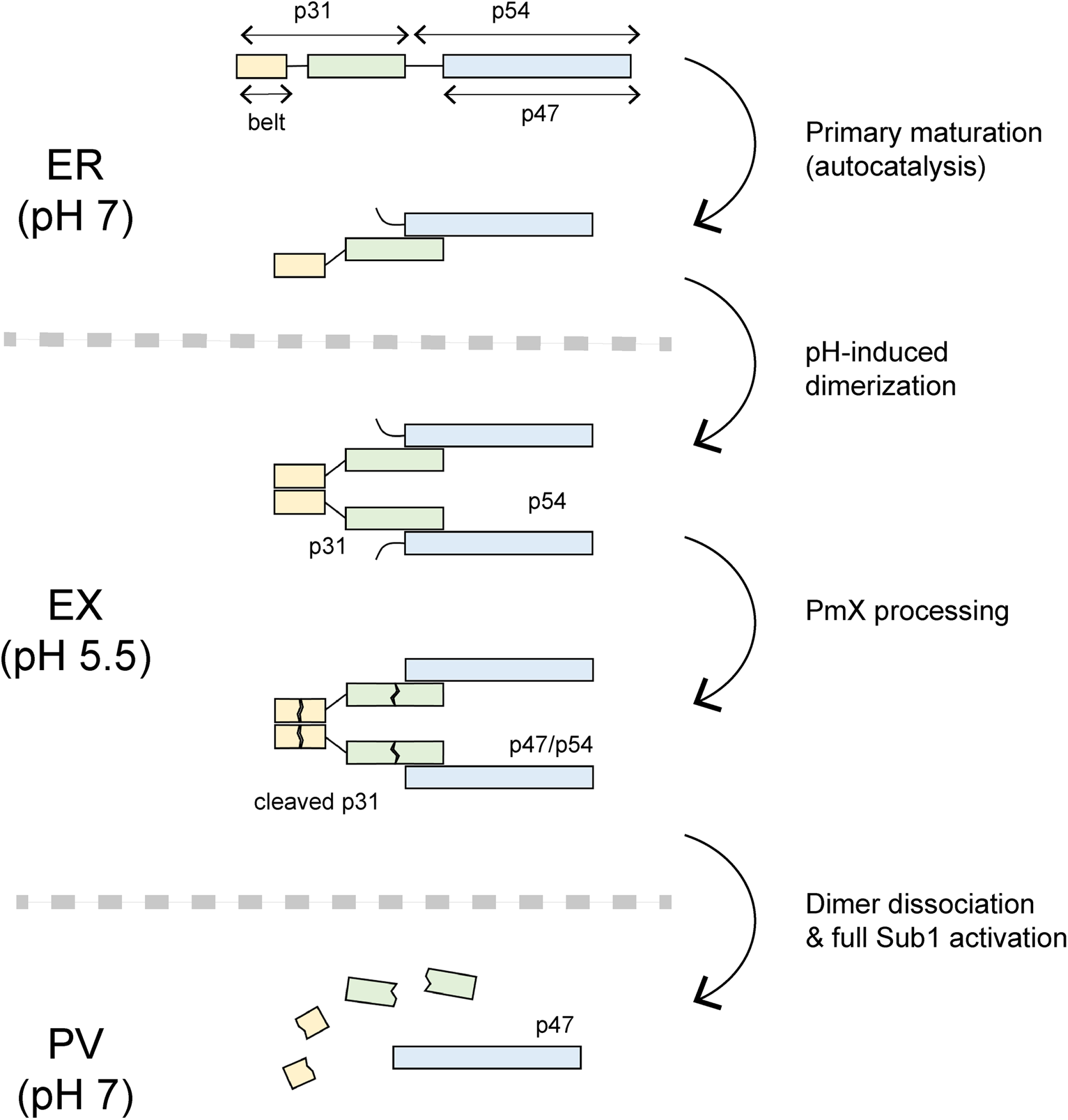
Model for the activation of Sub1 in *Plasmodium falciparum*. Primary maturation of Sub1 into the PD (p31) and the catalytic domain (p54) species takes place early in the endoplasmic reticulum (ER). The acidic environment of the exoneme (EX) induces the formation of the PD-mediated autoinhibited Sub1 dimer, which would remain inactive upon cleavage by plasmepsin X (PmX). Upon discharge into the rather neutral parasitophorous vacuole (PV), the higher pH dissociates the dimer leading to full Sub1 activation. The conversion of the catalytic domain from the p54 to the p47 species can be obtained either by direct PmX cleavage in the exoneme or by autocatalysis in the PV.

It has been recently reported that the aspartic protease PmX cleaves the Sub1 PD, as an intact form of PD (p31) could be detected in parasite lysates lacking PmX (44). In agreement with the acidic environment of secretory vesicles (53, 54), the exoneme-localized PmX requires acidic conditions for activity (42, 43, 55, 56), implying that Sub1 PD cleavage should occur during exoneme transport, before its discharge into the rather neutral PV (57), which is the site of Sub1 action (34, 58). In other words, PD processing and Sub1 activation appear to occur at different times in different cell compartments. The results reported here strongly suggest that this spatiotemporal dissociation is made possible by pH-dependent Sub1 dimerization. According to a plausible model (Figure 7), the dimeric assembly of Sub1 is maintained within the acidic environment of the exoneme, even after PmX processing, to preserve not only the structural integrity of the protease but also its complete inactivation during transport. This hypothesis is consistent with the crystal structures, as the two PmX cleavage sites detected in the PD, respectively in the middle of the N-terminal helix of the belt subunit and in a solvent-exposed loop from the bacterial PD-like subunit (44) are on the surface of the molecule and are not expected to interfere with dimer formation.

Upon the cGMP-mediated drastic shift in [Ca^2^] that immediately precedes the egress of merozoites (31), Sub1 is discharged into the PV (33) where the increase of pH would then promote dimer dissociation and the subsequent clearance of PD fragments (44), leading to full Sub1 activation. This last step within the PV may likely involve further Sub1 auto-catalytic processing of the PD, as two cleavage sites have been detected in the long loop from the PvS1 belt subunit in the presence of calcium (46). As for the catalytic domain, the conversion of the p54 into the p47 species at Asp249 (52) can also take place in both compartments, either by PmX in the exoneme or by auto-catalysis in the PV (44). Essentially, this N-terminal processing corresponds to cleave off the poorly conserved (46) unstructured region from the core subtilase domain (Figure 7) and was recently shown to be dispensable for parasite growth (44).

For the subtilase family in general, the displacement of PDs following primary automaturation is possibly the most important and rate-limiting single event for activation. These may involve variations of [Ca^2^] and/or pH, as described for bacterial subtilisins (12) or for eukaryotic furin and PC1/3, which undergo subtle pH-induced conformational changes that regulate their activation within secretory vesicles (16, 17). Variations in pH also participate in removing the PD of plant subtilisins (18) and a lower pH during secretion participates in activating yeast proteinase A and procarboxypeptidase Y (59, 60). For Sub1, *Plasmodium* appears to use a novel pH-dependent two-step strategy, as activation requires both an endogenous activating protease, PmX, to cleave the PD and a pH increase (arising from the discharge of Sub1 into a different cell compartment) to relieve inhibition. Although it is unclear why the malaria parasite does it this way, this novel regulatory element of subtilisin activation – so far only observed in *Plasmodium* – is part of a more complex process (involving several enzymes, cellular compartments and signaling events) required to ensure the egress of competent merozoites from RBCs and allow *Plasmodium* propagation in the host blood and subsequent vector borne transmission.

## Materials and methods

### Production and purification of the PfS1FL, PvS1FL, PvS1CD and PvS1βBelt recombinant enzymes

The baculovirus-expressed recombinant PfS1_FL_, PvS1_FL_ and PvS1β_Belt_ (Genebank accession codes FJ536585, JX491486 and KM211548, respectively) were prepared as described (46, 61, 62). Briefly, culture supernatant was harvested 3 days post-infection with recombinant baculoviruses, centrifuged 30 min at 2150*g* and concentrated/diafiltrated against loading buffer (D-PBS, 500 mM NaCl, 5 mM Imidazole) using the AKTA crossflow system (GE healthcare) supplemented with a 10KDa, 1.2ft2 Kvicklab cassette (GE healthcare).

The proteins were purified on an AKTA purifier system (GE Healthcare) at 4 °C. The sample was loaded onto a 5-ml TALON metal affinity resin (Clontech) equilibrated in loading buffer. After extensive washes with loading buffer, the bound protein was eluted with a linear gradient of 5–200 mM imidazole in D-PBS, 500 mM NaCl. Purified protein-containing fractions were pooled and concentrated by using Amicon Ultra 15 (molecular weight cutoff of 10,000) and size-fractionated onto a HiLoad 16/60 Superdex 75 column equilibrated with 20 mM Tris, 500 mM NaCl, 1 mM CaCl_2_ (pH 8). Fractions were monitored by absorbance (280 nm) and analyzed by Coomassie Blue staining of SDS-polyacrylamide gels and enzyme activity assay. For enzymatic assays, purified recombinant proteins PvS1_FL_ and PvS1 catalytic domain (PvS1_CD_), prepared as described in Martinez *et al.* (63) were stored at -20 °C following the addition of 10% v/v of pure glycerol. Protein concentrations were determined from A_280_, using the extinction coefficient predicted by ExPASy ProtParam (64).

### Protein crystallization

Initial identification of crystallization conditions was carried out using the vapor diffusion method in a Cartesian technology workstation. Sitting drops were set using 200 nl of 1:1 mixture of PfS1_FL_ and a crystallization solution (672 different conditions, commercially available), equilibrating against 150 μl reservoir in a Greiner plate. Optimization of initial hits was pursued manually in Linbro plates with a hanging drop setup. The best crystals were obtained by mixing 1.5 μl of native PfS1_FL_ (9.7 mg/mL) with 1.5 μl of the reservoir solution containing 1.6 M NaH_2_PO_4_ / 0.4 M K_2_HPO_4_ and 0.1 M phosphate-citrate, pH 5.2, at 18°C. Rod-like crystals appeared within 3-4 weeks and had dimensions of 0.25 x 0.05 x 0.05 mm. Prior to diffraction data collection, single crystals were flash-frozen in liquid nitrogen using a mixture of 50% paratone and 50% paraffin oil as cryoprotectant.

### Data collection, structure determination, and refinement

X-ray diffraction data were collected at 100 K using beamline Proxima1 (wavelength = 1.07169 Å) at the SOLEIL synchrotron (France). All datasets were processed using XDS (65) and AIMLESS from the CCP4 suite (66). The crystal structure was determined by molecular replacement methods using Phaser (67) and *P. vivax* Sub1 (PDB code 4tr2) as the probe model. Crystallographic refinement was done through iterative cycles of manual model building with COOT (68, 69) and reciprocal space refinement with BUSTER (70) using a TLS model and non-crystallographic symmetry restraints. The crystallographic statistics are shown in Table 1. Structural figures were generated with Chimera (71) or Pymol (The PyMOL Molecular Graphics System, Version 2.0 Schrödinger, LLC). Atomic coordinates and structure factors of PfS1_FL_ have been deposited in the protein data bank under the accession code 8POL.

### Analytical Ultracentrifugation (AUC)

Prior to AUC analysis, protein samples were dialyzed against buffer pH 8 (50 mM Tris pH 8, 150 mM NaCl, 10 mM CaCl2) or buffer pH 5 (50 mM NaCit pH 5, 150 mM NaCl, 10 mM CaCl2) at 4 °C during 24 hs. Protein samples were centrifuged in a Beckman Coulter XL-I analytical ultracentrifuge at 20 °C in a four-hole rotor AN60-Ti equipped with 12 mm 6 sectors centerpieces. Detection of the protein concentration as a function of radial position and time was performed by optical density measurements at 280 nm. For sedimentation equilibrium experiments, 120 μl of PvS1or PvS1β(2 μM, 5 μM, and 10 μM) in buffer pH 8 or buffer pH 5 were spun sequentially at rotor speeds of 9000 rpm, 13000, and 16000 rpm. The data were acquired after reaching equilibrium for one hour at each speed. The following parameters were calculated using Sednterp 1.09 and used for the analysis of the experiment: partial specific volume of 0.732ml g^−1^, viscosity *η* = 1.068 cP, and density *ρ* = 1.0139g ml^−1^.

Sedimentation equilibrium radial distributions were analyzed by global fitting using the single specie model with Ultrascan 9.5 software (http://www.ultrascan.uthscsa.edu/“\t”_blank) (72). Monte Carlo analysis was performed on the fit in order to estimate measurement error.

### PvS1 enzymatic activity

The activity of PvS1_FL_ and PvS1_CD_ was assessed as previously described, at 37°C for 60 min in 10 mM CaCl_2_, 150mM NaCl at different pH (61). Briefly, the reaction was initiated by adding 20 µM of a FRET substrate (Dabsyl-KLVGADDVSLA-EDANS) and followed by measuring the EDANS fluorescence (excitation at 360 nm and emission at 500 nm) every 2 min under shaking over 1 h using a Tecan Infinite M1000 spectrofluorimeter. All measurements were performed in triplicate.

### Culture of *P. falciparum* parasites, western blots and immunofluorescence analysis

Parasites were cultured as previously described (61) in RPMI 1640 medium containing L-glutamine, 25 mM HEPES (Invitrogen) supplemented with 2,5% Albumax II (Gibco), with a hematocrit of human red blood cells of 5%, 100 µM hypoxanthine (C.C.pro, Germany), 25 µg/ml gentamycin (SigmaAldrich) at 37 °C in a 5% O_2_, 5% CO_2_, and 90% N_2_ atmosphere. To obtain human red blood cells, human peripheral blood samples were collected from healthy volunteers through the ICAReB platform (Clinical Investigation & Access to Research Bioresources) from the Center for Translational Science, Institute Pasteur (73). All participants received an oral and written information about the research and gave written informed consent in the frame of the healthy volunteers Diagmicoll cohort (Clinical trials NCT 03912246) and CoSImmGEn cohort (Clinical trials NCT 03925272), after approval of the Ethics Committee of Ile-de-France (2009, April 30th and 2011, jan 18th, respectively).

*P.falciparum* schizonts, enriched for mainly mature forms prior to merozoite egress, were obtained as previsoulsy described (74) from a parasite culture enriched in schizonts on Percoll 75% prior to be incubated in standard culture conditions until the expected parasite stages are obtained. Following washes of the mature segmented schizonts in PBS, proteins were extracted in n-dodecyl-β-D-maltoside (DDM, Merck) 1% detergent, diluted in PBS supplemented with protease inhibitors (Complete, Roche), during 30 minutes on ice and centrifuged at 16100 g at 4°C for 15 minutes. After addition of native sample buffer (0.1% Ponceau S/50% glycerol), the protein extracts were separated on a NativePAGE 4-16% Bis- Tris using NativePAGE 20X Running Buffer (Invitrogen) for the anode buffer, supplemented with DOC (Na-Deoxycholate, Sigma) 0.05% and 0.01% DDM for the cathode buffer. After migration (80V/1h plus 150V/2h30), polyacrylamide gels were incubated 15min at room temperature with agitation in a denaturation buffer composed of transfer buffer (Biorad) supplemented with 2% SDS and 10mM DTT (Sigma). Proteins were then transferred on a nitrocellulose membrane using a Trans-Blot Turbo (Biorad). The NativeMark Unstained Protein Standard (Invitrogen) is revealed by incubation of the membrane in a 0.2% Ponceau S solution (Serva). For SDS page conditions, same samples were separated and transferred as previously described (61).

The full-length PfS1 recombinant purified protein was used to immunized rabbits subcutaneously following a two-months standard procedure (Proteogenix, France). Briefly, the first immunization by the purified protein emulsified with the complete Freund’s adjuvant was followed by two boosts with incomplete Freund’s adjuvant at days 39 and 53 and the serum were collected at day 70 post the initial injection and kept in 50% glycerol at -20 °C.

Blots were incubated with antibodies (1/3000 dilution) purified on G-protein (Proteogenix, France), followed by horseradish peroxidase-conjugated (HRP) secondary rabbit antibodies (Promega, 1/25000) and revealed by chemiluminescence (Pierce) using ChemiDoc and ImageLab (BioRad).

Immunofluorescence analysis were performed as previously described (61) using Percoll- purified *P. falciparum* segmented schizonts washed in PBS, deposited on glass slides and air- dried before to be fixed with paraformaldehyde 4% and permeabilized with 0.1% Triton X100 (Sigma). Slides were saturated with 1% Albumax II overnight at 4°C before successive incubations with the purified anti-PfS1 and the anti-MSP1 mouse mAb G17.12 both diluted to 1:1000. Secondary antibodies Alexa Fluor 488-conjugated anti-rabbit IgG and Alexa Fluor 594-conjugated anti-mouse IgG (Invitrogen) were used diluted to 1:1000. Antibodies were prepared in PBS containing Albumax II 1%. Hoescht 33342 (Invitrogen) diluted at 1:5000 was added to the secondary antibody solutions. PBS-washed slides were sealed with Vectashield (Vector laboratories). Images were collected using a Leica DM5000B microscope with a x1000 magnification.

## Acknowledgements.

We thank the staff of the Crystallography core facility at the Institut Pasteur for carrying out robot-driven crystallization screenings. We acknowledge ESRF for provision of synchrotron radiation facilities and we thank the staff of the Proxima 1 beamline for support during data collection. We are grateful to the healthy volunteers for their participation in the study and thanks the staff, particularly Hélène Laude et Emmanuel Roux, of ICAReB-Clin and ICAReB- biobank of the CRBIP (BioResource Center) from the Medical Direction of the Institute Pasteur for managing the visits of healthy volunteers and for preparing and providing the human blood samples used to cultivate *P.falciparum*.

